# The differential effect of age on upper limb sensory processing, proprioception and motor function

**DOI:** 10.1101/2022.08.26.505430

**Authors:** Leen Saenen, Geert Verheyden, Jean-Jacques Orban de Xivry

## Abstract

Sensory processing consists in the integration and interpretation of somatosensory information. It builds upon proprioception but is a distinct function requiring complex processing by the brain over time. Currently little is known about the effect of aging on sensory processing ability, nor the influence of other covariates such as motor function, proprioception, or cognition. In this study, we measured upper limb passive and active sensory processing, motor function, proprioception, and cognition in 40 healthy younger adults and 54 older adults. We analyzed age differences across all measures and evaluated the influence of covariates on sensory processing through regression. Our results showed larger effect sizes for age differences in sensory processing (r=0.38) compared to motor function (r=0.18-0.22) and proprioception (r=0.10-0.27), but smaller than for cognition (r=0.56-0.63). Aside from age, we found no evidence that sensory processing performance was related to motor function or proprioception, but active sensory processing was related to cognition (β=0.30-0.42). In conclusion, sensory processing showed an age-related decline, while some proprioceptive and motor abilities were preserved across age.

## Introduction

Intact somatosensory function is essential for normal daily life functioning (1). Through perception of external and internal information (i.e., exteroception and proprioception) and subsequent cortical processing (i.e., sensory processing) to interpret this information, the body is able to control motor actions (1, 2). In the context of aging, various studies have investigated the possible existence of an age-related decline in proprioceptive abilities (3–8), however, the effect of age on sensory processing has been less extensively researched. Here, proprioception is defined as the detection of limb position and movement coming from muscle spindle receptors and Golgi tendon organs by the primary somatosensory cortex (9), while sensory processing is defined as a higher cortical function which requires integration and interpretation of proprioceptive and exteroceptive information in the secondary somatosensory cortex in order to build concrete concepts, which makes it a distinct and more complex function (2, 10). For example, while proprioception provides information on shoulder, elbow and hand position when manipulating objects, sensory processing combines all information coming from different sources (i.e., exteroceptive and proprioceptive sources such as cutaneous receptors and muscle spindles (1, 2)) over time to recognize and name the object. Sensory processing has alternatively been described as tactile discrimination, proprioceptive discrimination, or haptic perception (11, 12). Somatosensory function may be impaired in neurological disorders, such as stroke, which can hamper motor recovery and negatively affect daily life functioning (13, 14). Interestingly, it has been shown that sensory processing can be affected after stroke even when proprioception and exteroception are intact (15), and that it shows a distinct relationship with daily life functioning (16), which highlights the importance of investigating sensory processing separately.

Recently, our group has developed a robot-based sensory processing assessment consisting of a passive and active condition (17). These novel assessments showed good discriminative and convergent validity to evaluate sensory processing function in participants with chronic stroke (17). Performance on these tasks may be influenced by different covariates such as age. Indeed, previous studies have reported a decline in clinical sensory processing abilities in healthy aging (18, 19). In addition, given that sensory processing is a secondary higher cortical function which processes primary exteroceptive and proprioceptive information (2), proprioceptive abilities may influence outcomes on the sensory processing assessments. Building further on this assumption, proprioception may also be affected by age (3, 19), although some authors did not report this finding (4–8). Furthermore, motor function and cognition may influence task performance, given that the task requires active movement and storage of information in the working memory. Both motor function and cognition may also be affected by age (20, 21). In summary, it is important to identify which factors influence performance, in order to interpret scores on the sensory processing assessments.

The aim of this study was (1) to evaluate differences in sensory processing abilities between healthy younger and older adults, as well as on other assessments of sensorimotor and cognitive function; (2) to assess the relationship between the outcomes of the sensory processing tasks and motor functions, proprioception, and cognition while taking age into account; and (3) to also re-evaluate the influence of age on the robot-based sensory processing assessments while considering possible variability in motor function, proprioception, and cognition. We hypothesized older adults would perform worse on the sensory processing tasks than younger adults, and that these results would be comparable to other assessments of sensorimotor and cognitive function. In addition, we hypothesized that proprioception and cognition would be related to both the passive and active conditions (given the processing of proprioceptive information, and the reliance on working memory, respectively, for both conditions), and that they would reduce the effect of age.

## Material and Methods

### Participants

Forty healthy younger adults and 54 healthy older adults participated in this cross-sectional study. Younger adults were aged between 18 and 30 years old, while the older adults were at least 55 years old. Participants were excluded when they had history of musculoskeletal or neurological disorders (such as stroke), or presented with upper limb sensorimotor impairments. The group of older participants was also included in a previous study (17). All participants provided written informed consent before participation. This study was approved by the ethical committee UZ/KU Leuven (S61997) and was registered at clinicaltrials.gov (NCT04723212).

### Experimental set-up

The Kinarm End-Point robot (BKIN Technologies Ltd., Kingston, Canada) was used for all robot-based assessments. This bimanual robot allows passive and active movement in the horizontal plane through grasping of the end-point handles and uses a virtual reality screen to provide control of visual feedback of the upper limbs. An additional black cloth was used to ensure there was no remaining vision of the upper limbs. All assessments were performed seated.

### Experimental task

A novel sensory processing task was used, which has been described in detail elsewhere (17). Both the passive and the active conditions of the task were performed, and they consisted of exploration, reproduction, and identification of geometrical shapes (Fig. 1A and 1B). In the passive condition, the robot first passively moved the participant’s non-dominant arm in the shape of a triangle, tetragon, or pentagon, after which the participant was asked to actively reproduce the shape without mirroring with the dominant arm. In the active condition, the participant was asked to first explore the same shapes by moving the non-dominant arm between virtual walls which delimited the shape, and then to reproduce the shape with the dominant arm. In both conditions, participants were finally asked to identify the explored shape out of six options presented on the robot screen, and to indicate how certain they were of this answer with a 4-point Likert scale. The task consisted of 15 randomized shapes which were preceded by 5 practice trials. Visual feedback of the upper limbs was blocked during the task, and feedback on task performance was only provided during the practice trials. In the passive condition, a bell-shaped speed profile was used with a maximum speed of 0.67 m/s. In the active condition, position-dependent force regions were used to delimit the shape. Along the lines of the shape, the participant could actively move within a 0.2 cm wide zero force region. Outside these lines, the robot applied virtual walls with a stiffness of 6000 N/m and viscosity of –50 Ns/m. Participants could explore each shape once within a time limit of 30 seconds. Five parameters were calculated from this task using custom MATLAB (MathWorks, Natick, USA) scripts (17):

**Figure 1.**
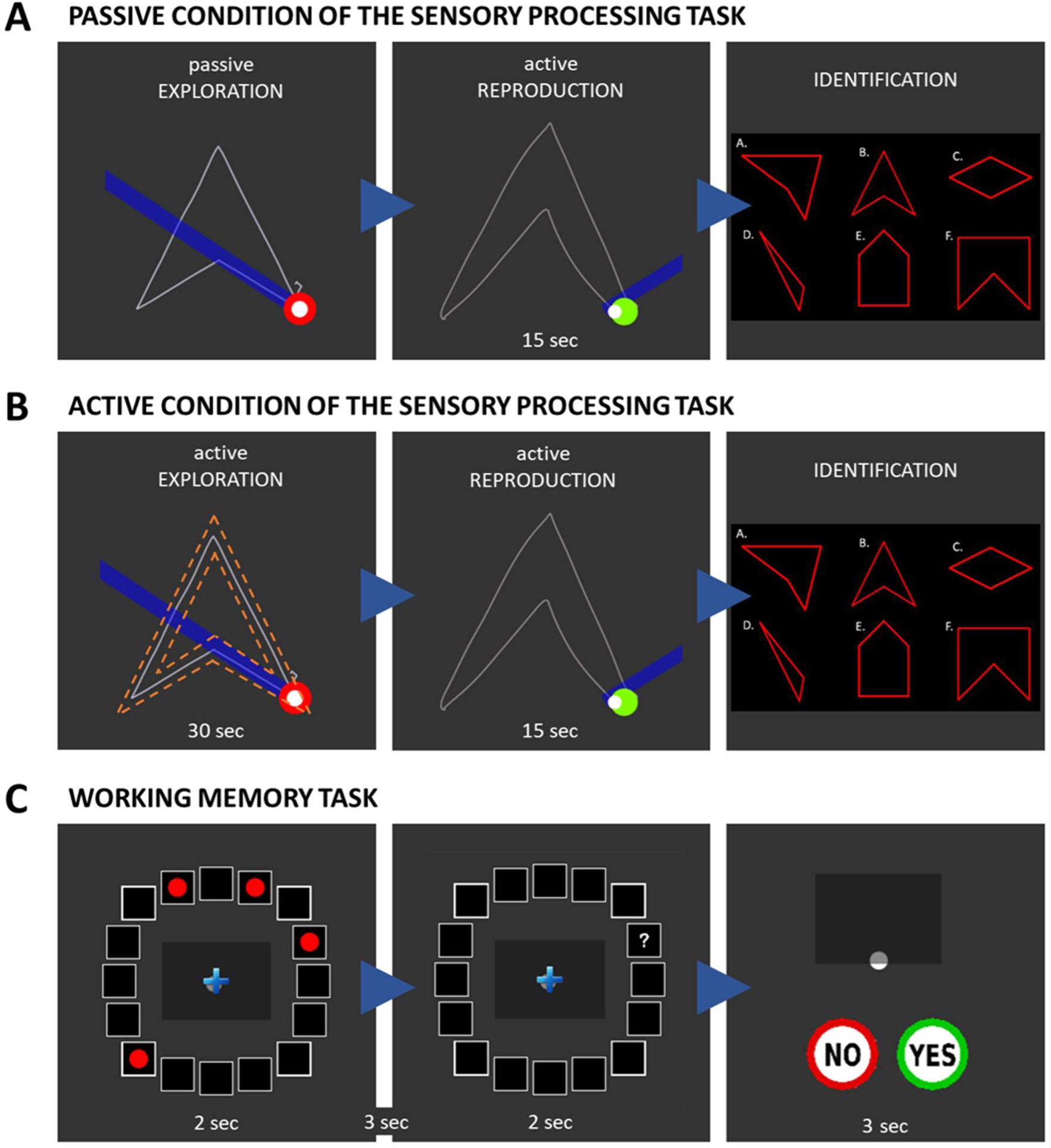
A Passive condition of the sensory processing task. Left panel: Passive exploration of the shape with the non-dominant arm. Middle panel: Reproduction of the shape with the dominant arm. Right panel: Identification of the explored shape. **B Active condition of the sensory processing task**. Left panel: Active exploration of the shape with the non-dominant arm. Middle panel: Reproduction of the shape with the dominant arm. Right panel: Identification of the explored shape. White solid line = arm movement path. Orange dashed lines = invisible virtual walls delimiting the shape. **C Working memory task**. Left panel: Three, four, five or six targets are shown. Middle panel: Question mark appears in or close to one of the target locations. Right panel: Responding whether a target was present in the indicated location.

- Cross-correlation X and Y: mean similarity between explored and reproduced shapes by cross-correlating hand position signals on the X-and Y-axes; values range between –1 and 1
- Dynamic time warping: mean similarity between explored and reproduced shapes by optimally aligning the two temporal sequences of hand position signals (i.e., minimizing the sum of the Euclidean distances between corresponding points by stretching one temporal sequence against the other); values equal the sum of the Euclidean distances between all corresponding points of the two aligned sequences (in m)
- Procrustes analysis: mean similarity between explored and reproduced shapes by optimally superimposing both shapes through translation, rotation and scaling of the reproduced shape on top of the explored shape; values range from 0 to 1, and equal the standardized sum of the Euclidean distances between corresponding points of both superimposed shapes
- % correctly identified: the percentage of correctly identified shapes

### Other robot-based assessments

A visually guided reaching task was performed bilaterally to assess motor function (22). In this task, participants were asked to perform reaching movements to 4 targets centered around a central target. Ten outcome parameters quantifying feedforward and feedback control were calculated across 20 trials for each arm using Dexterit-E software (BKIN Technologies Ltd., Kingston, Canada), including posture speed, reaction time, initial direction angle, initial distance ratio, speed maxima count, min-max speed, movement time, path length ratio, and max speed. A detailed description of the outcome parameters is found elsewhere (22).

In addition, an arm position matching task was performed bilaterally to assess proprioception (23). In this task, the robot first moved the participant’s arm to one of 9 targets, after which the participant was asked to mirror-match this position with the other arm. The task is performed without visual feedback of both arms. The following parameters were reported for each arm across 54 trials (23):

- Absolute error XY: mean absolute unsigned distance error (in m) by calculating the root-sum-square of absolute errors on the X-and Y-axis
- Variability XY: mean variability in signed hand position (in m) by calculating the root-sum-square of standard deviations on the X-and Y-axis

Finally, a working memory task was performed to assess cognition (Fig. 1C) (24). Here, 16 squares were positioned in a circle on the robot screen. Participants were asked to remember the positions of three, four, five or six targets which simultaneously lit up for 2 seconds within the squares. Then, after a 3 second delay, a question mark appeared for 2 seconds in or close to one of the target positions. Participants indicated whether a target had appeared in the indicated location by moving their dominant arm to “Yes” or “No” within 3 seconds. The task consisted of 48 trials, and we report two outcome parameters:

- Total score: number of correct answers
- Capacity: estimation of the number of targets which can be stored in the working memory. The following formula was used: K = S*(H-F), where K is the working memory capacity, S the number of targets, H the hit rate (i.e., number of correct answers divided by total number of trials) and F the false alarm rate (i.e., number of wrong answers divided by total number of trials) (24–26). A more detailed description of the calculation can be found in Supplemental Figure S1.

### Clinical assessments

The SENSe Assess tool was used to quantitatively evaluate upper limb somatosensory function (27). The following three assessments were performed:

- Wrist position sense test (28): assessment of wrist proprioception in which the examiner passively flexed or extended the participant’s wrist, after which the participant was asked to move a pointer to the perceived wrist position. The average error (in degrees) between actual and indicated position was calculated based on 20 trials.
- Tactile discrimination test (29): assessment of sensory processing through discrimination of different sets of finely graded textures. The participant was asked to explore three texture surfaces of which two were identical, and to indicate which one was different. Outcomes include the total number of correct answers out of 25 trials, and the area under the curve.
- Functional tactile object recognition test (30): assessment of sensory processing through identification of different everyday objects using touch only. Since a ceiling effect was observed in this study for the number of correct answers, we only reported the average time needed to explore the objects across 14 trials.

Abstract shape representation was assessed with the shape drawing test of the Montreal cognitive assessment (31). A score of 1 was assigned in case of a correctly executed drawing of a three-dimensional cube, whereas a score of 0 was assigned in case of incorrect drawing.

### Statistical analysis

All statistical analyses were performed in R version 4.2.0 (32). Statistical tests were performed two-tailed with an alpha level of 0.050. Participant characteristics were described using medians, interquartile ranges, and percentages, and compared between the younger and older participants with Mann-Whitney U tests and Fisher’s Exact tests (‘wilcox.test’ and ‘fisher.test’ from the stats package (32), respectively).

We used factor analysis to assess the underlying factors that explain the relationships among a set of observed variables obtained in some tasks where many outcomes are reported (sensory processing and visually-guided reaching tasks). Therefore, the exploratory factor analyses were used to reduce our dataset by combining the parameters of the sensory processing tasks and the visually guided reaching task into one variable/factor per task. Factor extraction was performed with the principal factor extraction method (33) on age-standardized data for the passive condition of the sensory processing task, active condition of the sensory processing task, and the visually guided reaching test separately (‘fa’ from the psych package (34)). Factor extraction was based on scree plots and Kaiser’s criterion (i.e., eigenvalue > 1), after confirmation of adequate sample size with the Kaiser-Meyer-Olkin test and appropriate correlation coefficients between variables (i.e., most correlations being between 0.30 and 0.90) (35). In case of more than one factor, ‘oblimin’ rotation was used. This way, parameters which are correlated with each other are assumed to represent to same underlying construct. Factor loadings were obtained during the factor extraction step, which are the correlations between each parameter and the factor (35). Factor scores were finally obtained for each test using the original unstandardized data with the regression method (‘factor.scores’ from the psych package (34)). The reported factor scores are z-scores, so they can be interpreted as the deviation from average performance. Positive factor scores represent better than average performance, while negative factor scores mean worse than average performance.

Next, we evaluated differences between younger and older adults on cross-correlation values of the sensory processing tasks using a robust three-way ANOVA (‘bwwtrim’ from Wilcox 2017), with age group (younger vs. older adults) as between-group factor, and task condition (passive vs. active) and axis direction (X vs. Y) as within-group factors. For dynamic time warping values, Procrustes values, the percentage of identified shapes, and certainty values, a robust two-way ANOVA (‘bwtrim’ from Wilcox 2017) was calculated with participant group as between-group factor and task condition as within-group factor. For all ANOVA analyses, generalized eta squared (η^2^ η^2^G) was calculated to indicate effect size (‘anova_summary’ from the rstatix package (37)) (38, 39). Next, age differences for the factor scores of the passive and active sensory processing tasks, as well as all other assessments were evaluated using Mann-Whitney U tests, because Shapiro-Wilk tests indicated a non-normal distribution for most outcomes. For each test, the effect size r was calculated (40). We calculated 95% confidence intervals for the effect size r by bootstrapping 1000 samples (‘boot’ and ‘boot.ci’ from the boot package (41, 42)).

Then, the relationship between the passive and active sensory processing tasks, and motor function, proprioception and cognition was evaluated using robust moderated multiple regression analyses (‘lmRob’ from the robust package (43)). The factor scores of the sensory processing tasks were estimated by including the outcome of interest (i.e., motor function, proprioception, or cognition), the age group (young=-1 vs. old=1), and their interaction as independent variables. All scores were first converted into z-scores to obtain standardized regression coefficients. Here, we report the standardized regression coefficient β of the robot-based and clinical assessments to indicate their relationship with the sensory processing tasks regardless of age. 95% confidence intervals of the regression coefficients were obtained using the following formula: β ± 1.96 * SE, with SE being the standard error. Note that the results for the shape drawing test of the Montreal cognitive assessment are only narrative described given the crude dichotomous scoring.

Finally, results from the above-described robust moderated multiple regression analyses were also used to re-evaluate the age differences on the factor scores of the passive and active sensory processing tasks, while considering possible influence of variability in motor function, proprioception, and cognition. For this purpose, the *p*-value associated with the regression coefficient of age was inspected.

## Results

Forty healthy younger adults and 54 healthy older adults underwent evaluation of sensory processing, motor function, and proprioception. Forty younger adults and 40 older adults also underwent additional evaluation of cognition. Participant characteristics can be found in Table 1. There was an equal distribution of gender and hand dominance between both age groups, however, the younger adults received a median of one extra year of education.

**Table 1.** Participant characteristics.

|  | Younger adults<br>(*n* = 40) | Older adults<br>(*n* = 54) | *P*-value<sup>†</sup> | Older adults<br>Working memory task<br>(*n* = 40) | *P*-value<sup>‡</sup> |
| --- | --- | --- | --- | --- | --- |
| Median age in years (IQR) | 24.24 (22.73-25.16) | 62.89 (57.63-67.95) | < 0.001* | 62.39 (57.43-67.19) | < 0.001* |
| Gender, *n* (%) |  |  | 0.41 |  | 0.50 |
| Male | 15 (38) | 25 (46) |  | 19 (48) |  |
| Female | 25 (63) | 29 (54) |  | 21 (53) |  |
| Hand dominance, *n* (%) |  |  | 0.16 |  | 0.26 |
| Right | 34 (85) | 51 (94) |  | 38 (95) |  |
| Left | 6 (15) | 3 (6) |  | 2 (5) |  |
| Median years of education (IQR) | 17 (16-18) | 16 (15-18) | 0.02* | 16 (15-18) | 0.06 |
| Level of education, *n* (%) |  |  | 0.17 |  | 0.11 |
| Primary education | 0 (0) | 0 (0) |  | 0 (0) |  |
| Lower secondary education | 0 (0) | 3 (6) |  | 1 (3) |  |
| Higher secondary education | 6 (15) | 10 (19) |  | 6 (15) |  |
| Bachelor's degree | 11 (28) | 21 (39) |  | 19 (48) |  |
| Master's degree | 23 (58) | 20 (37) |  | 14 (35) |  |

\* $p < 0.050$ <sup>†</sup> Comparison between older adults and younger adults with Mann-Whitney U tests and Fisher's Exact tests.<sup>‡</sup> Comparison between older adults who completed the working memory task and younger adults with Mann-Whitney U tests and Fisher's Exact tests.

### Older adults were less accurate in reproducing explored shapes than younger adults

To quantify reproduction of the passive and active sensory processing tasks, cross-correlation values were calculated on the X-and Y-axes. We found that older adults showed significantly lower cross-correlation values (mean 0.85, SD 0.05) than younger adults (mean 0.88, SD 0.04), meaning that older adults did not reproduce the shape as accurately as younger adults (Fig. 2A; main effect of age: F(1,86) = 23.13, *p* < 0.001, η^2^G = 0.14). Cross-correlation values were also significantly worse for the active condition (mean 0.84, SD 0.05) than for the passive condition (mean 0.88, SD 0.04; Fig. 2A; main effect of condition: F(1,86) = 96.52, *p* < 0.001, η^2^G = 0.25). We found no evidence that group differences differed across the two conditions (Fig. 2A; age x condition: F(1,86) = 0.29, *p* = 0.59, η^2^G < 0.01). We also did not find evidence for any other two-way interactions (Fig. 2A; age x direction: F(1,86) = 0.03, *p* = 0.86, η^2^G < 0.01; condition x direction: F(1,86) = 1.87, *p* = 0.17, η^2^G < 0.01), or that cross-correlation values were different on the X-or Y-axis (Fig. 2A; main effect of direction: F(1,86) < 0.01, *p* = 0.95, η^2^G < 0.01). We did find a significant three-way interaction (Fig. 2A; age x condition x direction: F(1,86) = 5.62, p = 0.02, η^2^G < 0.01), but this was not further explored given the small effect size and the fact that the difference between axis directions was not considered relevant here.

**Figure 2.**
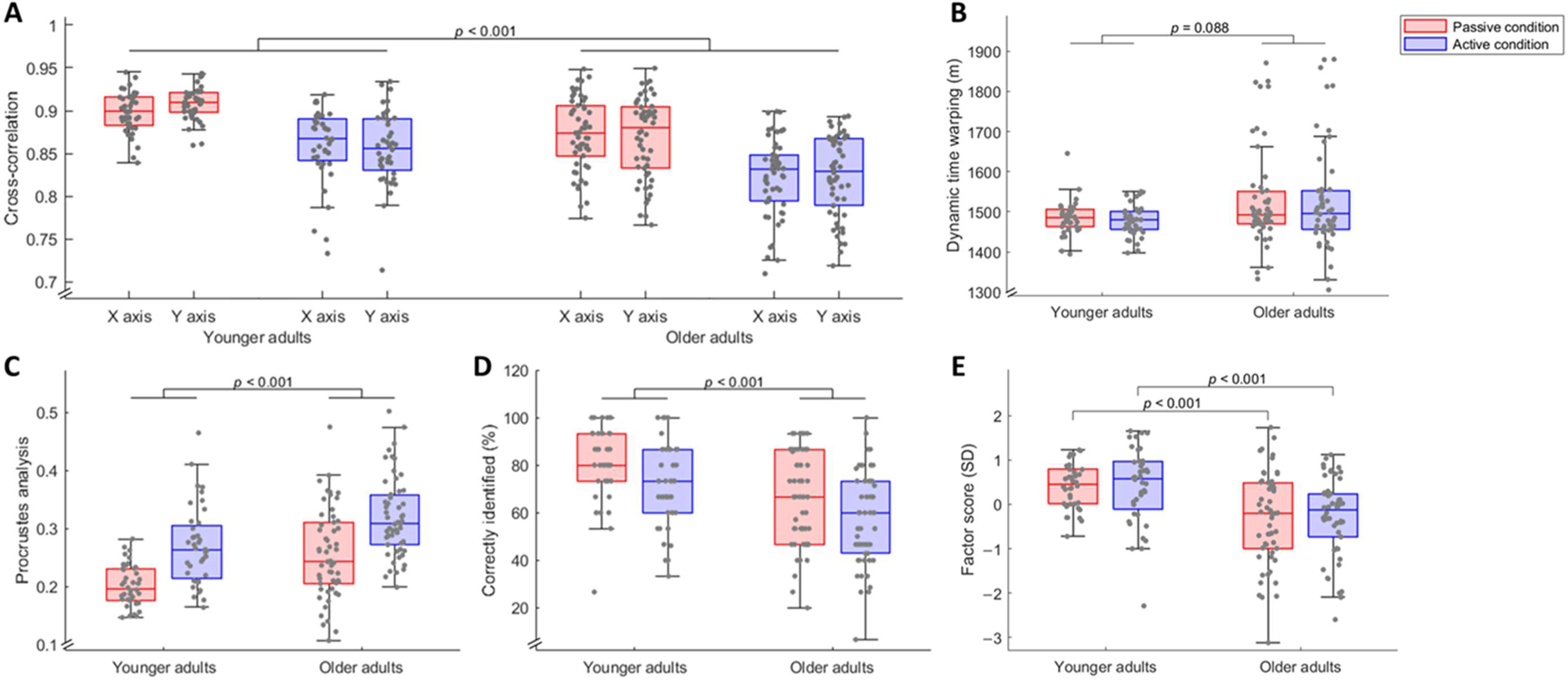
Results of the passive (in red) and active (in blue) sensory processing assessments. Visualization of boxplot with 25^th^, 50^th^ and 75^th^ percentile (box) indicated and largest and lowest values within 1.5 times the interquartile range (error bars). Dots represent individual participant results. **A** Main effect for between-group analysis of three-way ANOVA for cross-correlation on X and Y axes. Higher values are associated with better performance. **B** Main effect for between-group analysis of two-way ANOVA for dynamic time warping. Lower values are associated with better performance. **C** Main effect for between-group analysis of two-way ANOVA for Procrustes analysis. Lower values are associated with better performance. **D** Main effect for between-group analysis of two-way ANOVA for the percentage of correctly identified shapes. Higher values are associated with better performance. **E** Between-group comparison of factor scores using Mann-Whitney U tests. Higher values are associated with better performance.

In addition to the cross-correlation values, dynamic time warping values and Procrustes values were calculated to evaluate the shapes as one entity. For dynamic time warping, we found no evidence of a significant difference between older adults (mean 1530, SD 126) and younger adults (mean 1481, SD 39; Fig. 2B; main effect of age: F(1,90) = 3.04, *p* = 0.09, η^2^G = 0.06). Dynamic time warping values were also not significantly different between the active condition (mean 1507, SD 105) than the passive condition (mean 1511, SD 98; Fig. 2B; main effect of condition: F(1,90) = 0.85, *p* = 0.36, η^2^G < 0.01), and we found no interaction between age group and task condition (Fig. 2B; age x condition: F(1,90) = 0.46, p = 0.50, η^2^G < 0.01). Similar results were found for the Procrustes analysis, as older adults showed significantly worse scores (mean 0.29, SD 0.08) than younger adults (mean 0.24, SD 0.06; Fig. 2C; main effect of age: F(1,90) = 21.79, *p* < 0.001, η^2^G = 0.13), and values were also worse for the active condition (mean 0.30, SD 0.07) than the passive condition (mean 0.23, SD 0.07; Fig. 2C; main effect of condition; F(1,90) = 52.40, *p* < 0.001, η^2^G = 0.20). Again, we did not find evidence that performance in each task condition differed as a function of age (Fig. 2C; age x condition; F(1,90) = 0.01, *p* = 0.92, η^2^G < 0.01).

### Older adults identified less shapes correctly, despite being equally certain about their answers, than younger adults

During the identification phase, older adults (mean 62.01, SD 20.11) identified significantly less shapes correctly than younger adults (mean 76.24, SD 16.85; Fig. 2D; main effect of age: F(1,90) = 16.93, *p* < 0.001, η^2^G = 0.13). Identification was also worse for the active condition (mean 64.12, SD 19.87) than for the passive condition (mean 72.02, SD 19.50; Fig. 2D; main effect of condition: F(1,90) = 16.58, *p* < 0.001, η^2^G = 0.05), but age differences did not differ across task conditions (Fig. 2D; age x condition: F(1,90) = 0.45, *p* = 0.50, η^2^G < 0.01). Certainty about the answers did not differ between both groups (younger adults: mean 2.27, SD 0.48; older adults: mean 2.14, SD 0.47; main effect of age: F(1,90) = 2.04, *p* = 0.16, η^2^G = 0.02), but was significantly lower for the active condition than the passive condition (main effect of condition: F(1,90) = 14.93, *p* < 0.001, η^2^G = 0.04). There was a mean certainty of 2.12 (SD 0.51) for the active condition, and 2.28 (SD 0.43) for the passive condition, which indicates high certainty for both conditions. We found no evidence of a two-way interaction effect (age x condition: F(1,90) = 0.59, *p* = 0.45, η^2^G < 0.01).

### Older adults have reduced sensory processing ability based on the novel assessment

To sum up our results, we performed a factor analysis with the parameters of the sensory processing tasks for the active and passive conditions separately. Dynamic time warping was excluded from this factor analysis because it was not correlated with the other parameters. Scree plots indicated that a single factor was present for both the passive and the active condition of the sensory processing task. Factor loadings for each parameter can be found in the Supplemental Table S1. The factor scores of the passive and active sensory processing tasks give an indication of the overall sensory processing ability, taking into account reproduction and identification parameters. For the passive condition, we found that older adults had a median factor score of –0.20 (IQR –0.99-0.49), while younger adults showed a significantly higher median of 0.46 (IQR 0.01-0.79; Fig. 2E; U = 598, *p* < 0.001, r = 0.38). For the active task, we also found a significant difference between older adults (median –0.13, IQR –0.71-0.24) and younger adults (median 0.58, IQR –0.06-0.96; Fig. 2E; U = 596, *p* < 0.001, r = 0.38). We found medium effect sizes for both conditions (Fig. 4).

### Older adults have lower working memory and clinical sensory processing abilities than younger adults, but maintain some proprioceptive and motor abilities

We entered all parameters of the visually guided reaching test into a factor analysis. Three of them (hand posture speed, reaction time and initial speed ratio) were excluded from this analysis because they did not show correlations with the other parameters. Two factors were extracted, representing motor control and speed, respectively. The factor loadings for each parameter can be found in the Supplemental Table S1. Between-group analysis on both factors scores showed that older adults had worse motor control and lower speed than younger adults, but only small effect sizes were found (Fig. 3 and 4; motor control: U = 852, *p* = 0.082, r = 0.18; speed: U = 801; *p* = 0.03; r = 0.22). For the arm position matching test, we found a significant difference for absolute error XY of the dominant arm, and variability XY of the non-dominant arm, but again, only small effect sizes were found (Fig. 3 and 4; absolute error XY dominant arm: U = 1384, <u>p</u> = 0.010, r = 0.24; absolute error XY non-dominant arm: U = 1232, *p* = 0.25, r = 0.12; variability XY dominant arm: U = 1201, *p* = 0.25, r = 0.10; variability XY non-dominant arm: U = 1416, *p* = 0.01, r = 0.27). Older adults did not show a reduced wrist position sense (Fig. 3; U = 1069, *p* = 0.93, r = 0.01), while they did show reduced sensory processing abilities as assessed with the tactile discrimination test (Fig. 3; total score: U = 714, *p* = 0.005, r = 0.29; area under curve: U = 716, *p* = 0.005, r = 0.29) and functional tactile object recognition test (Fig. 3; U = 1576, *p* < 0.001, r = 0.39). Finally, older adults also showed a significantly lower working memory than younger adults, with large effect sizes (Fig. 3 and 4; total score: U = 276, *p* < 0.001, r = 0.63; capacity: U = 366, *p* < 0.001, r = 0.56).

**Figure 3.**
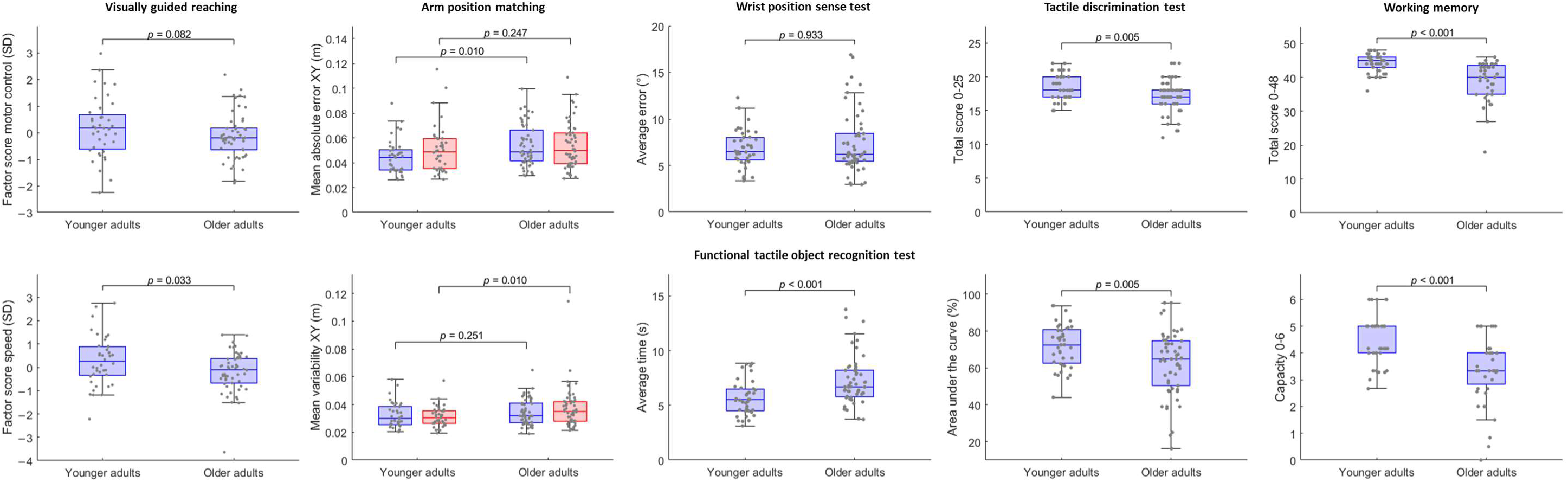
Between-group comparisons using Mann-Whitney U tests for the factor scores of the visually guided reaching test, mean absolute error XY and variability XY of the arm position matching test (dominant arm in blue, non-dominant arm in red), average error of the wrist position sense test, average time of the functional tactile object recognition test, total score and area under the curve of the tactile discrimination test, and working memory total score and capacity. Visualization of boxplot with 25^th^, 50^th^ and 75^th^ percentile (box) indicated and largest and lowest values within 1.5 times the interquartile range (error bars). Dots represent individual participant results.

In summary, while large and medium effect sizes were found for cognition and sensory processing, respectively, we only found small effect sizes for proprioception and motor function (Fig. 4).

**Figure 4.**
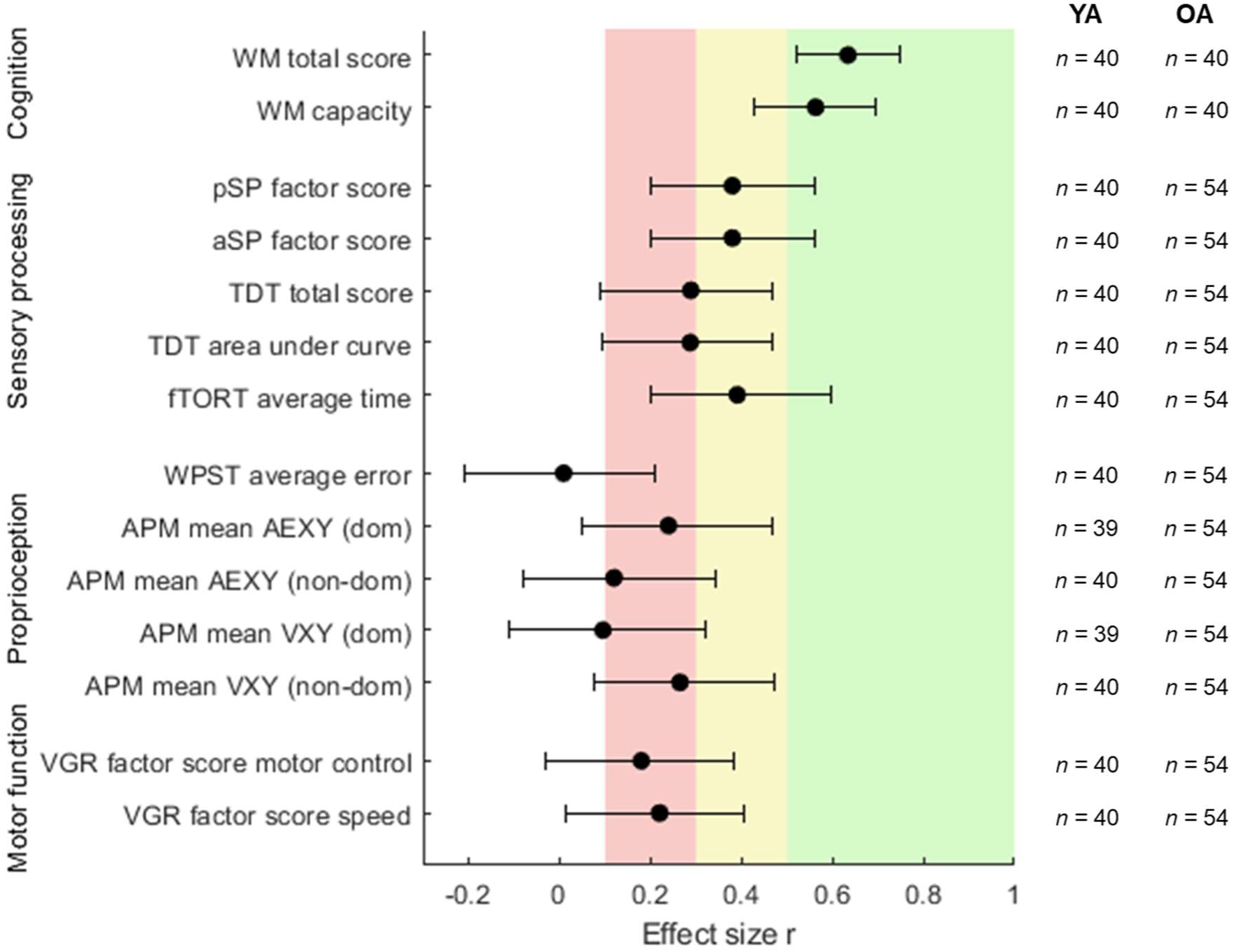
Comparison of effect sizes and 95% confidence intervals of age differences between assessments of cognition, sensory processing, proprioception, and motor function. Red = small effect size; Yellow = medium effect size; Green = large effect size. *Abbreviations:* YA = younger adults; OA = older adults; WM = working memory; pSP = passive condition of sensory processing task; aSP = active condition of sensory processing task; TDT = tactile discrimination test; fTORT = functional tactile object recognition test; WPST = wrist position sense test; APM = arm position matching; AEXY = absolute error XY; dom = dominant arm; non-dom = non-dominant arm; VXY = variability XY; VGR = visually guided reaching

### Working memory is related to the factor score of the active sensory processing task

For the active condition, the working memory total score and capacity were found to have moderate relationships with the factor score regardless of age (Table 2). Motor function and proprioception showed overall only weak associations with the sensory processing assessment.

**Table 2.** Results of regression equation Y = β_0_ + β_1_.X_1_ + β_2_.AGE + β_3_.X_1_.AGE. Y is the factor score of the sensory processing task, X_1_ the outcome of interest, and AGE the dichotomized age group. β_1_ indicates the relationship between the outcome of interest and the sensory processing assessment regardless of age. The *p*-value associated with AGE indicates whether there is still a significant relationship between age and the sensory processing assessment when taking the outcome of interest into account. The *p*-value associated with β_3_ indicates whether an interaction effect exists between the outcome of interest and age.

| **PASSIVE CONDITION OF THE SENSORY PROCESSING TASK** |  |  |  |  |  |  |  |  |  |
| --- | --- | --- | --- | --- | --- | --- | --- | --- | --- |
| $X_1$ | $\beta_1$ | [95% CI] | $P$-value | AGE | [95% CI] | $P$-value | $\beta_3$ | [95% CI] | $P$-value |
| Visually guided reaching |  |  |  |  |  |  |  |  |  |
| Factor score motor control | -0.19 | [-0.45 0.06] | 0.14 | -0.37 | [-0.62 -0.12] | 0.005* | -0.15 | [-0.40 0.10] | 0.25 |
| Factor score speed | 0.07 | [-0.16 0.31] | 0.55 | -0.35 | [-0.59 -0.11] | 0.005* | 0.01 | [-0.23 0.25] | 0.94 |
| Arm position matching |  |  |  |  |  |  |  |  |  |
| Absolute error XY |  |  |  |  |  |  |  |  |  |
| Dominant arm | -0.25 | [-0.51 0.01] | 0.06 | -0.28 | [-0.52 -0.03] | 0.03* | -0.16 | [-0.42 0.09] | 0.22 |
| Non-dominant arm | -0.21 | [-0.48 0.06] | 0.13 | -0.35 | [-0.61 -0.08] | 0.01* | -0.14 | [-0.41 0.13] | 0.31 |
| Variability XY |  |  |  |  |  |  |  |  |  |
| Dominant arm | -0.27 | [-0.52 -0.01] | 0.04* | -0.31 | [-0.57 -0.06] | 0.02* | -0.11 | [-0.36 0.15] | 0.42 |
| Non-dominant arm | -0.19 | [-0.54 0.16] | 0.30 | -0.32 | [-0.54 -0.09] | 0.01* | 0.05 | [-0.30 0.40] | 0.80 |
| Working memory |  |  |  |  |  |  |  |  |  |
| Total score | 0.13 | [-0.20 0.47] | 0.45 | -0.37 | [-0.65 -0.09] | 0.01* | 0.20 | [-0.13 0.54] | 0.25 |
| Capacity | 0.13 | [-0.14 0.39] | 0.35 | -0.38 | [-0.64 -0.12] | 0.005* | 0.18 | [-0.08 0.44] | 0.18 |
| Wrist position sense test |  |  |  |  |  |  |  |  |  |
| Average error | 0.09 | [-0.18 0.36] | 0.52 | -0.38 | [-0.59 -0.17] | 0.001* | 0.00 | [-0.27 0.27] | 0.99 |
| Tactile discrimination test |  |  |  |  |  |  |  |  |  |
| Total score | 0.06 | [-0.20 0.31] | 0.66 | -0.35 | [-0.59 -0.11] | 0.006* | 0.02 | [-0.23 0.28] | 0.85 |
| Area under curve | 0.14 | [-0.13 0.40] | 0.33 | -0.32 | [-0.56 -0.09] | 0.009* | 0.10 | [-0.17 0.36] | 0.49 |
| Functional tactile object recognition test |  |  |  |  |  |  |  |  |  |
| Average time | -0.08 | [-0.38 0.23] | 0.63 | -0.34 | [-0.60 -0.09] | 0.01* | -0.18 | [-0.49 0.13] | 0.26 |

| **ACTIVE CONDITION OF THE SENSORY PROCESSING TASK** |  |  |  |  |  |  |  |  |  |
| --- | --- | --- | --- | --- | --- | --- | --- | --- | --- |
| $X_1$ | $\beta_1$ | [95% CI] | *P*-value | AGE | [95% CI] | *P*-value | $\beta_3$ | [95% CI] | *P*-value |
| Visually guided reaching |  |  |  |  |  |  |  |  |  |
| Factor score motor control | −0.19 | [−0.43 0.05] | 0.12 | −0.41 | [−0.64 −0.19] | 0.001* | −0.21 | [−0.45 0.03] | 0.09 |
| Factor score speed | 0.02 | [−0.27 0.31] | 0.89 | −0.35 | [−0.64 −0.06] | 0.02* | −0.19 | [−0.47 0.10] | 0.20 |
| Arm position matching |  |  |  |  |  |  |  |  |  |
| Absolute error XY |  |  |  |  |  |  |  |  |  |
| Dominant arm | −0.05 | [−0.28 0.18] | 0.67 | −0.36 | [−0.59 −0.14] | 0.002* | −0.12 | [−0.35 0.11] | 0.31 |
| Non-dominant arm | −0.19 | [−0.43 0.24] | 0.60 | −0.36 | [−0.70 −0.02] | 0.04* | −0.07 | [−0.40 0.27] | 0.69 |
| Variability XY |  |  |  |  |  |  |  |  |  |
| Dominant arm | −0.31 | [−0.52 −0.11] | 0.004* | −0.35 | [−0.55 −0.15] | 0.001* | 0.03 | [−0.18 0.23] | 0.80 |
| Non-dominant arm | −0.25 | [−0.66 0.16] | 0.24 | −0.32 | [−0.61 −0.04] | 0.03* | −0.05 | [−0.46 0.36] | 0.83 |
| Working memory |  |  |  |  |  |  |  |  |  |
| Total score | 0.42 | [−0.20 1.05] | 0.19 | −0.25 | [−0.74 0.23] | 0.31 | 0.08 | [−0.55 0.70] | 0.81 |
| Capacity | 0.30 | [−0.03 0.64] | 0.08 | −0.29 | [−0.61 0.04] | 0.09 | −0.02 | [−0.35 0.32] | 0.92 |
| Wrist position sense test |  |  |  |  |  |  |  |  |  |
| Average error | −0.12 | [−0.43 0.19] | 0.45 | −0.35 | [−0.61 −0.10] | 0.008* | 0.12 | [−0.19 0.43] | 0.44 |
| Tactile discrimination test |  |  |  |  |  |  |  |  |  |
| Total score | 0.03 | [−0.23 0.29] | 0.84 | −0.35 | [−0.61 −0.10] | 0.007* | 0.10 | [−0.17 0.36] | 0.47 |
| Area under curve | 0.05 | [−0.20 0.31] | 0.67 | −0.36 | [−0.58 −0.13] | 0.003* | 0.15 | [−0.10 0.40] | 0.25 |
| Functional tactile object recognition test |  |  |  |  |  |  |  |  |  |
| Average time | −0.04 | [−0.35 0.28] | 0.82 | −0.36 | [−0.64 −0.07] | 0.02* | −0.03 | [−0.34 0.28] | 0.85 |

\* $p < 0.050$ . Abbreviations: $\beta$ = standardized regression coefficient; CI = confidence interval

Seven older adults did not succeed in the shape drawing test of the Montreal cognitive assessment, and their results on the passive and active conditions were variable (passive condition: factor score –1.78-0.52; active condition: factor score –2.09-0.26).

### Working memory influences age differences on the active sensory processing task

Even when considering the influence of variability in motor function, proprioception, and cognition, the effect of age on the passive sensory processing task remained significant (Table 2), meaning that age is an important contributor to variability in performance on the passive sensory processing task. For the active condition, however, the addition of working memory reduced the significant role of age (Table 2). We did not find evidence of a significant interaction between age and any of the other outcome measures (Table 2).

## Discussion

In this study, 94 healthy younger and older adults performed novel passive and active sensory processing assessments. We found that older adults have reduced upper limb sensory processing ability compared to younger adults, as seen by less accurate reproduction and identification of geometrical shapes in the absence of visual feedback. Interestingly, we found medium effect sizes for age differences in sensory processing ability, while only small effect sizes were found for motor function and proprioception. For cognition, large effect sizes were found. In fact, working memory showed a moderate relationship with the active condition of the sensory processing assessment, and it also reduced the significant role of age. For the passive condition, we found that age was the largest contributor to variability in performance, and that neither motor function, nor proprioception, nor cognition were substantially related to performance on this task.

The current findings are in line with results of previous studies investigating sensory processing ability across age (18, 19). In the study of Master et al. (2010), younger adults scored on average 91.1% during a tactile letter recognition task, while older adults identified 79.2% on average (18). While our task was possibly more difficult, given the lower identification scores in both age groups, the difference between younger and older adults was similar (14.2 in the present study vs. 11.9 in the study of Master et al. (2010)). Please note that very similar age groups were used in both studies. In our study, we found a larger effect size for sensory processing than for proprioception, confirming that they should be regarded as distinct modalities. In fact, the role of age on proprioception is currently under debate. We found only small differences for some of the parameters of the arm position matching task. Some studies found no differences in the same task for the dominant arm (7, 8), in contrast to another study who did suggest an age-related decline (3). It has been argued that only physically inactive older adults show decreased upper limb proprioception, while active older adults do not (5, 6, 44). No data on physical activity is available for the present study, but it’s likely the older adults in this study were physically active as they were recruited from a university sports center. Similarly, we found smaller differences across age for motor function than what was previously assumed (21). Yet, our findings are in line with another recent study which found only small effect sizes of age on motor function (45).

We can only speculate about the origin of the difference in age-related deterioration of sensory processing compared to proprioception. A possible explanation for this differential effect of age on sensory processing and proprioception could be linked to cortical activation patterns. While the detection of proprioceptive information is mainly located in the primary somatosensory cortex, sensory processing activates additional brain areas such as the secondary somatosensory cortex (46). Therefore, sensory processing requires activation of a broader cortical network as compared to proprioception. Age-related changes have been found in both the primary and the secondary somatosensory cortex (47, 48) and throughout the sensorimotor network (49). If sensory processing requires more cortical processing, then it could be more heavily affected by dedifferentiation of brain activation patterns (neurons respond to a larger range of stimuli (here a larger range of position information) with age (50, 51)), by increased noise (the signal-to-noise ratio is reduced in elderly people compared to young people (52)), or by the reduced number of neurons recruited by a given task (loss of functionality of synapses (53)). Alternatively, it is also possible that age-related changes do not occur uniform across the brain (54) and that the primary somatosensory cortex is less affected than brain areas involved in sensory processing, which might explain the differential effect of age on sensory processing, proprioception, and motor function. Further research is needed on the evaluation of sensory processing, proprioception, and motor function across age.

We have aimed to develop a quantitative assessment of upper limb sensory processing, without substantial influence of other confounding factors (17). In the present study, we investigated the influence of motor function, proprioception and cognition on the passive and active conditions of the sensory processing assessment. We did not find evidence that performance on the passive condition showed a relationship with motor function, proprioception, or cognition, apart from a small relationship with variability XY of the dominant arm during an arm position matching task. We hypothesized we would find a larger relationship with proprioception, given that the task relies on sensory processing of mainly proprioceptive information. However, the current results emphasize that sensory processing is nonetheless a distinct function, as was previously proposed by others (2, 55). Results differ for the active condition, where we did find a moderate positive relationship with working memory, meaning reduced working memory was associated with reduced performance on the active condition of the sensory processing assessment. In contrast, abstract shape representation did not seem to be related to the passive and active conditions, but these results should be interpreted with caution given the dichotomous scoring. Interestingly, the moderate relationship with working memory suggests that the active condition is a more complex task to perform in comparison to the passive condition. In fact, the active condition may be regarded as a dual task, where participants are required to combine active motor planning with creating a mental image of the shape, whereas in the passive condition, a larger focus can be placed on the latter. The results imply that attention should be paid to the cognitive functioning of participants when performing the active condition. Nonetheless, the results presented here suggest that the described evaluation protocol provides an accurate representation of upper limb sensory processing.

Some limitations of the study should be acknowledged. First, our results showed large confidence intervals for the presented effect sizes and regression coefficients, therefore, further studies are needed to confirm our results with more certainty. Second, most included participants received higher education, which might have led to high cognitive functioning in both age groups. Consequently, the relationship with cognition might have been underestimated. Finally, we have attempted to map several upper limb functions with use of standardized assessments. However, it should be acknowledged that the assessments are not perfect and only capture certain aspects of upper limb function. For instance, it is unclear whether visually-guided reaching captures all the inter-subject variability in motor function that could influence performance at the sensory processing task. Likewise, several somatosensory assessments require transferring information from one arm to the other, which could modulate our measure of somatosensory function but is not related to that concept.

## Conclusions

Sensory processing is a distinct function from proprioception, as the latter is defined as the detection of limb position and movement, whereas the former requires additional complex integration of somatosensory information in order to interpret stimuli. We found that there is a medium decline in sensory processing abilities across age, while proprioception and motor function show only a small decline. In fact, we found that sensory processing is mostly related to age and less by proprioception or motor function. Cognition might be an additional confounder when assessing active sensory processing.

## Acknowledgements

The authors would like to acknowledge Arthur Booms and Joppe Groffils for their assistance with recruitment and data collection. We are grateful to all participants in this study.

## Grants

This work was supported by KU Leuven Internal Funds [grant number C22/18/008].

## Disclosures

none

## Notes

### Competing Interest Statement

The authors have declared no competing interest.

### Summary of Updates

Results corrected; Figure 2 corrected; Figure 3 revised; Figure 4 corrected; Table 2 corrected; Table S1 corrected; Introduction clarified; Outcome measure added; Methods clarified

