## Supplemental Figure S1 for "The differential effect of age on upper limb sensory processing, proprioception and motor function"

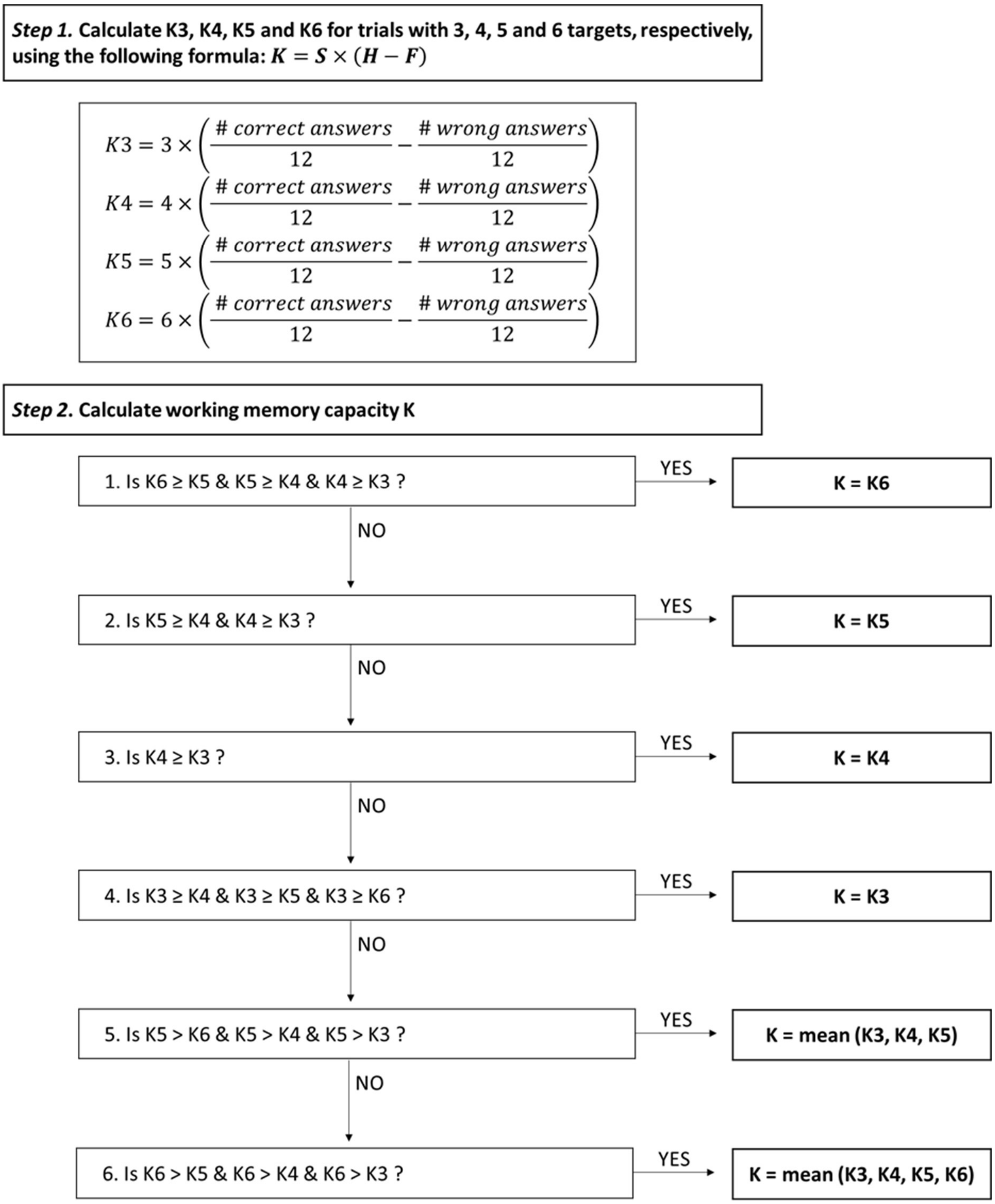


**Figure S1.** Calculation of the working memory capacity.

*Abbreviations*: K = working memory capacity; S = size of array; H = hit rate; F = false alarm rate.
