## Supplemental Table S1 for "The differential effect of age on upper limb sensory processing, proprioception and motor function"

**Table S1.** Factor loadings of the parameters of the active and passive condition of the sensory processing tasks, and the visually guided reaching task.

| | Sensory processing | |
|---|---|---|
| | Passive condition | Active condition |
| Cross-correlation X | 0.84 | 0.83 |
| Cross-correlation Y | 0.82 | 0.87 |
| Procrustes analysis | -0.92 | -0.93 |
| % identified | 0.52 | 0.60 |
| | Visually guided reaching | |
| | Motor control | Speed |
| Mean initial direction angle | | |
| Dominant arm | 0.47 | 0.31 |
| Non-dominant arm | 0.52 | 0.14 |
| Mean initial distance ratio | | |
| Dominant arm | -0.84 | -0.01 |
| Non-dominant arm | -0.84 | 0.26 |
| Mean speed maxima count | | |
| Dominant arm | 0.63 | -0.18 |
| Non-dominant arm | 0.74 | -0.28 |
| Mean min-max speed | | |
| Dominant arm | 0.67 | 0.37 |
| Non-dominant arm | 0.63 | 0.40 |
| Mean movement time | | |
| Dominant arm | 0.11 | -0.89 |
| Non-dominant arm | 0.09 | -0.93 |
| Mean path length ratio | | |
| Dominant arm | 0.68 | 0.30 |
| Non-dominant arm | 0.67 | 0.32 |
| Mean maximum speed | | |
| Dominant arm | 0.34 | 0.74 |
| Non-dominant arm | 0.39 | 0.70 |
